# Whole genome base-wise aggregation and functional prediction for human non-coding regulatory variants

**DOI:** 10.1101/583237

**Authors:** Shijie Zhang, Yukun He, Huanhuan Liu, Haoyu Zhai, Dandan Huang, Xianfu Yi, Xiaobao Dong, Zhao Wang, Ke Zhao, Yao Zhou, Jianhua Wang, Hongcheng Yao, Hang Xu, Zhenglu Yang, Pak Chung Sham, Kexin Chen, Mulin Jun Li

## Abstract

Predicting the functional or pathogenic regulatory variants in the human non-coding genome facilitates the interpretation of disease causation. While numerous prediction methods are available, their performance is inconsistent or restricted to specific tasks, which raises the demand of developing comprehensive integration for those methods. Here, we compile whole genome base-wise aggregations, regBase, that incorporate largest prediction scores. Building on different assumptions of causality, we train three composite models to score functional, pathogenic and cancer driver non-coding regulatory variants respectively. We demonstrate the superior and stable performance of our models using independent benchmarks and show great success to fine-map causal regulatory variants. We believe that regBase database together with three composite models will be useful in different areas of human genetic studies, such as annotation-based casual variant fine-mapping, pathogenic variant discovery as well as cancer driver mutation identification. regBase is freely available at https://github.com/mulinlab/regBase.

## INTRODUCTION

Accurate prediction and prioritization of non-coding regulatory variants are crucial issues in the human genetic studies. Genome-wide association studies (GWASs) have produced numerous single-nucleotide variants (SNVs) that are associated with hundreds of medical traits and diseases, and the majority of the associations are suggested to be mediated by non-coding regulatory codes (1–3). Whole genome sequencing technologies are frequently incorporated into the relevance investigation of non-coding variants in Mendelian disease (4,5), and existing evidence also suggests that non-coding regulatory variants can modulate disease risk by affecting pathogenic coding variant penetrance (6). Given the high volume of disease-causal candidate variants in the regulatory region as well as the expensive downstream functional validations, computationally predicting non-coding regulatory variants has become important and long-standing scientific issue.

In the last few years, a large number of computational methods had been proposed to annotate and predict functional non-coding variants. Building on different predictive assumptions, abundant annotation datasets as well as complementary statistical models, these algorithms have achieved great successes to prioritize functional, pathogenic and cancer-relevant non-coding regulatory variants (7–10). However, the state-of-the-art benchmarks showed poor concordance among the prediction scores of several existing methods (11–13). To comprehensively evaluate the regulatory potential or pathogenesis of certain SNV outside the protein-coding region, researchers now have to collect and compare scores from different resources, even need to download huge pre-computed files or manually calculate prediction scores. The overwhelming growth of new prediction tools further complicates such retrieval processes. In addition, the incomplete understanding and the functional complexity of regulatory DNA impede the development of single but versatile model that is able to accurately predict causal regulatory variants affecting different biological processes. For example, recent commonly adopted algorithms that integrate evolutionary constraint, epigenomics, and sequence features, such as CADD (14,15), GWAVA (16), FunSeq2 (17) and fitCons (18), usually achieved limited predictive power for expression-modulating variants from *in vivo* saturation mutagenesis of an enhancer (19), or allele imbalanced variants influence critical molecular traits in the transcriptional regulation, like chromatin accessibility (20). Furthermore, compared with the functional regulatory variants prioritization, it is more challenging to predict pathogenic regulatory variants that underlie the development of Mendelian disorders or cancers (5,21). The insufficient accumulation of known pathogenic regulatory variants largely inhibits the characterization of their key discriminative features that is different from disease-free regulatory mutations.

In this work, we comprehensively integrate non-coding variant prediction scores from 23 tools for base-wise annotation of human genome, called regBase. As such, regBase provides first-time convenience to prioritize functional regulatory SNVs and assist the fine mapping of causal regulatory SNVs without queries from numerus resources. Inspired by the evident significance of ensemble prediction for pathogenic/deleterious nonsynonymous substitution, we systematically construct three composite models to score functional, pathogenic and cancer driver non-coding regulatory SNVs. We illustrate the discriminatory abilities and applicable scenarios of the proposed models by independent datasets and case study. regBase and associated models are freely available for download at https://github.com/mulinlab/regBase.

## MATERIALS AND METHODS

### Collecting, processing and integrating functional scores for non-coding regulatory variants

We downloaded base-wise precomputed scores for almost all possible substitutions of single nucleotide variant (SNV) in the human reference genome from 13 existing tools, including CADD (14,15), CDTS (22), CScape (23), DANN (24), Eigen (25), FATHMM-MKL (26), FATHMM-XF (27), FIRE (28), fitCons (18), FunSeq2 (17), GenoCanyon (29), LINSIGHT (30) and ReMM (31). We called this aggregated resource as regBase. For tool score recorded by interval-level value, such as CDTS, fitCons and LINSIGHT, we transformed continuous position into base-wise position and assigned the same score. Since some tools only support functional annotations for 1000 Genomes Project variants (32) or are inefficient to compute variant scores, we collected or generated functional scores of additional 10 tools for only biallelic variants from 1000 Genomes Project phase 3, including Basset (33), CATO (20), DanQ (34), DeepSEA (35), deltaSVM (36), FunSeq (37), GWAS3D (38), GWAVA_TSS (16), RSVP (39) and SuRFR (40) (see Table 1 and Supplementary Table S1 for details). We extracted 1000 Genomes Project biallelic variants from 13 base-wise precomputed scores and merged together with above 10 scores to generate a database that contains 23 tools for all biallelic variants, called regBase Common. Missing score values were replaced with “.” and genomic position of all variants were based on GRCh37/hg19. We also ranked all scores in each set and normalized them by PHRED-scaled score (−10*log10(rank/total)). The integrated database is tab delimited and indexed by Tabix (41).

### Correlation analysis

Three benchmark datasets were incorporated to evaluate the prediction consistency of existing tools including 1) the Human Gene Mutation Database (HGMD) functional regulatory variants used by GWAVA (42); 2) the ClinVar (201812 release) regulatory variants (43) with “CLNSIG=Pathogenic or CLNSIG=Benign” and only obtaining noncoding attributes by VEP (44) (not including splicing-altered consequences); 3) expression-modulating variants identified by massively parallel reporter assay (MPRA) with more than 1.5 log2 fold expression level change between alleles (45). Pearson correlation test and hierarchical clustering were used to evaluate the relationships of integrated tools upon these non-coding regulatory variant datasets, in which variants with missing value for any tools will be excluded.

### Construction of training dataset

We designed three training datasets to predict different categories of functional non-coding regulatory variants as follows:

regBase_REG dataset: functional regulatory variants regardless of functional direction and pathogenicity. We used our previously compiled functional regulatory variants dataset in PRVCS (11), which integrates four different resources including (i) the HGMD public dataset used by GWAVA; (ii) the ClinVar pathogenic variants in the non-coding region compiled by GWAVA; (iii) validated regulatory variants from the OregAnno database (46); (iv) fine-mapped disease-causal regulatory SNPs for 39 immune and non-immune diseases (47). Negative controls were sampled from allele frequency matched non-coding variants in the independent linkage disequilibrium (LD) with positive variants from 1000 Genomes Project.
regBase_PAT dataset: pathogenic regulatory variants. We incorporated ClinVar (201812 release) pathogenic regulatory mutations with “CLNSIG=Pathogenic” and only kept the mutations in the non-coding region by VEP annotations (not including splicing-altered consequences). We also included regulatory Mendelian mutations in the non-coding region from Genomiser (31) and merged with ClinVar data. For negative dataset, we randomly drew benign mutations with “CLNSIG=Benign” from ClinVar, and used the same strategy to retain non-coding mutations.
regBase_CAN dataset: cancer recurrent regulatory somatic mutations. For positive set, we downloaded COSMIC v84 non-coding mutations and selected ones having recurrence rate >= 10. For negative set, we sampled private non-coding somatic mutations with recurrence = 1 and PhyloP = 0 (48).

### Gradient Tree Boosting model

We made use of Gradient Tree Boosting (GTB) algorithm in our predictive model. In general, GTB is a special form of Gradient Boosting Machine, which makes prediction by combining the results of multiple weak learners, typically decision tree. We used XGBoost classifier as the implementation of GTB algorithm. XGBoost is a scalable end-to-end tree boosting system and has achieved the state-of-art performance in a large amount of tasks (49). Its sparsity-aware split finding makes it suitable for the task as missing value was commonly appeared in our datasets. We performed grid search based on 10-fold cross validation on training set in order to tune the hyper-parameters. While tuning training datasets with the unbalanced positive and negative samples, we adjusted the weight of positive samples according to the ratio of two classes. Receiver operating characteristic (ROC) curve and area under the receiver operating characteristics curve (AUC) were used to evaluate the performance of model during grid search.

### Construction of independent testing datasets

We assembled six independent testing datasets that were not used to train almost all of existing tools and our combined models, including 1) Brown_eQTL dataset: 11 tissue/cell type-specific eQTLs fine-mapping data that was profiled by Brown and colleagues (50). To further acquire more significant eQTL SNPs, we applied log10BF cutoff values of 10% FDR for each tissue/cell type; 2) GTEx_eQTL dataset: GTEx V6 44 tissues-specific eQTLs within CAVIAR (51) 95% fine-mapped credible set from UCSC (52); 3) MPRA_eQTL dataset: significant expression modulating variants (log2FC > 1.5) by MPRA in lymphoblastoid cell lines (45); 4) GWAS_5E-8 dataset: GWAS disease-associated variants with *P*-value < 5E-8 from GWAS Catalog v1.0.1 (53); 5) GWAS_1E-5 dataset: GWAS disease-associated variants with *P*-value < 1E-5 from GWAS Catalog v1.0.1 (53); 6) Somatic_eQTL dataset: recurrent somatic mutations from COSMIC V84 with recurrence >= 2 within significant flanking intervals per somatic eGene (54). We also generated corresponding controls for above datasets using different sampling strategies. For Brown_eQTL and GTEx_eQTL dataset, we randomly sampled allele frequency matched non-coding variants in the 10Kb transcription start site (TSS) regions of randomly selected genes. For GWAS_5E-8 and GWAS_1E-5 dataset, we sampled allele frequency matched non-coding variants in the independent LD with positive variants from 1000 Genomes Project. For MPRA_eQTL dataset, we used nonexpression-modulating variants (log2FC <0.005) by MPRA in lymphoblastoid cell lines. For Somatic_eQTL dataset, we sampled private non-coding somatic mutations from COSMIC V84 with recurrence = 1 and PhyloP = 0. We also excluded all positive and negative samples that have been incorporated in our training datasets.

### Evaluation schemes

We compared our composite models with integrated tools and two existing ensemble methods (PRVCS (11) and IW-Scoring (12)) using above six independent testing datasets. Positive predictive values (PPV), negative predictive values (NPV), false positive rate (FPR), false negative rate (FNR), sensitivity, specificity, accuracy, precision, recall, F1 score and Matthews correlation coefficient (MCC) were calculated according to Maximal Youden’s index during the measurement of ROC and AUC. We also calculated the correlation between true labels and prediction scores for each evaluation using Pearson correlation test.

### Causal variants prioritization for 5p15.33 TERT region

We collected significant trait/disease associated SNPs from GWAS catalog (*P*-value < 5E-8) and GWAS fine-mapping results from literatures at the 5p15.33 TERT region (Human GRCh37, chr5:1.22-1.37mb). We used LocusZoom (55) to visualize these disease-associated and fine-mapped SNPs on 1000 Genomes EUR population. To investigate the performance of regBase composite methods for causal variant prioritization, we extracted and normalized the raw scores of all tools in the 5p15.33 TERT region to generate regional PHRED-scaled scores. We further evaluated the sum or distribution of PHRED scores for all collected fine-mapped SNPs across different tools. Since some tools contain equal scores at this region and this will reduce the discrimination of true causal variants, we removed tools that obtain more than 25% equal scores in the evaluation.

## RESULTS

### Base-wise aggregation of non-coding regulatory variant prediction scores

We processed and compiled an integrative resource for prediction scores from 23 different tools on functional annotation of non-coding variants, including Basset (33), CADD (14,15), CATO (20), CDTS (22), CScape (23), DANN (24), DanQ (34), DeepSEA (35), deltaSVM (36), Eigen (25), FATHMM-MKL (26), FATHMM-XF (27), FIRE (28), fitCons (18), FunSeq (37), FunSeq2 (17), GenoCanyon (29), GWAS3D (38), GWAVA (16), LINSIGHT (30), ReMM (31), RSVP (39) and SuRFR (40) (Supplementary Table S1). Since some tools only support annotations for 1000 Genomes Project variants (32), or take long runtime to compute functional scores, we first built a database, called regBase Common, which contains functional scores from 23 tools for 38,248,779 in the 1000 Genomes Project phase 3. Among these integrated datasets, 13 tools provide precomputed scores for almost all possible substitutions of SNV in the human reference genome. Therefore, we also constructed a complete base-wise aggregation of non-coding variant functional scores for 8,575,894,770 substitutions of SNV, called regBase (Supplementary Table S2). We summarized the missing values in our integrated resources, and found that most of tools had less than 2% missing values across the whole genome. However, CATO (65.88%), SuRFR (33.91%) and CDTS (9.64%) exhibited relatively high or moderate missing rates in the regBase Common, and CDTS (13.02%) showed moderate missing rate in the regBase (Supplementary Table S3 and S4). To facilitate the efficient retrieve and comparison of functional scores of different alleles across tools, we indexed the whole dataset and used a PHRED-scaled method to normalize the raw score of each tool. The regBase and regBase Common can be downloaded from https://github.com/mulinlab/regBase.

### Correlation analysis of existing algorithms

Existing non-coding variants prediction algorithms dealt with different predictive objectives and assumptions, which could lead to inconsistent prediction on various application scenarios. To comprehensively evaluate the predictive concordance among our collected scores, we prepared three benchmark datasets that incorporate different pathogenicity/regulatory causality assumptions of non-coding regulatory variants (Supplementary Table S5): 1) functional regulatory variants from the public Human Gene Mutation Database (HGMD) (42) used by GWAVA; 2) pathogenic and benign regulatory variants from the ClinVar database (43); 3) experimentally validated expression quantitative trait loci (eQTL) variants from a massively parallel reporter assay (MPRA) (45). Pearson correlation analysis of regBase Common integrated functional scores showed both shared and distinct patterns on these benchmark datasets (Figure 1A). Algorithms trained on similar positive/negative data and features had relatively high pairwise correlations, like DeepSEA and DanQ (Pearson correlation coefficients R > 0.7), or CADD and DANN (R > 0.6), or FunSeq and FunSeq2 (R > 0.5). However, the majority of tools exhibited weak pairwise correlations (R < 0.4) in these regulatory variant datasets, which could be explained by the different training data and features, as well as the various learning models used. Among these tested noncoding regulatory variant datasets, we found the overall pairwise correlation for MPRA dataset was generally higher than those from other two datasets, implying that current tools may obtain better concordance in eQTL-associated regulatory variant prediction. Since some tested variants were not incorporated or obtained missing values in the regBase Common database, we also performed correlation analysis on 13 complete scores in the regBase database and found similar correlation patterns (Supplementary Figure S1).

**Figure 1.**
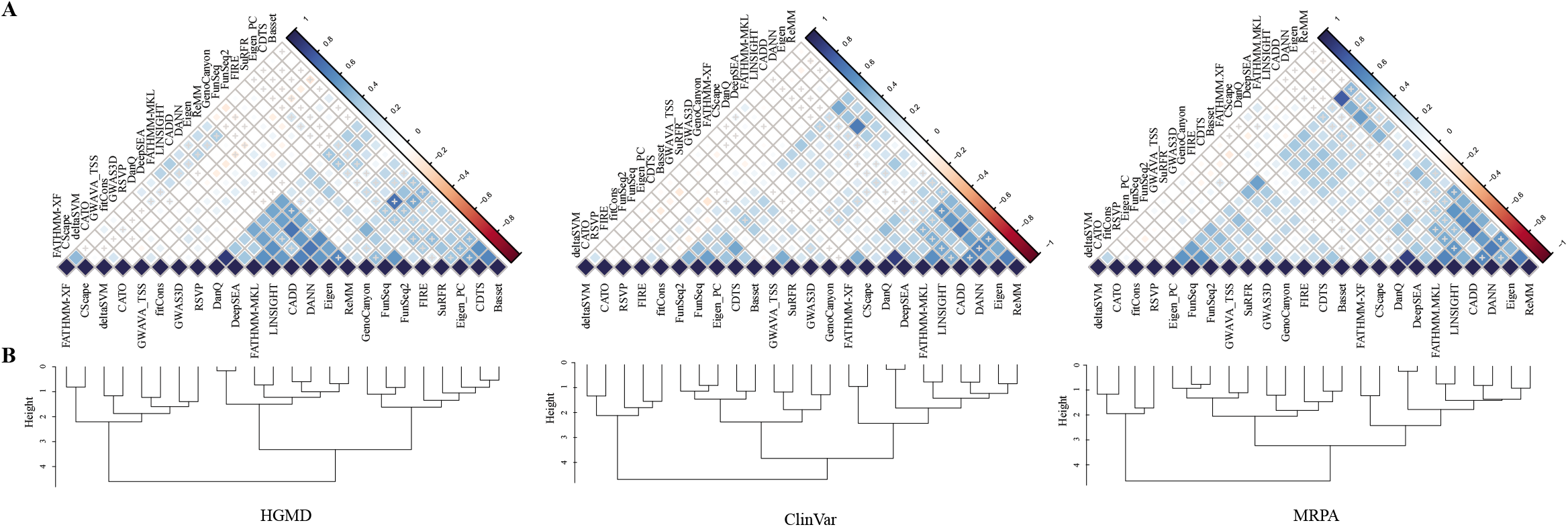
Correlation analysis of prediction score among 23 regBase Common integrated tools. (A) Pearson correlation of 23 regBase Common integrated functional scores on three known functional/pathogenic regulatory variant datasets. Positive correlations are displayed in blue and negative correlations in red color. Color intensity and the size of the square are proportional to the correlation coefficients. Nonsignificant *P*-value (>0.05) is marked with a cross. (B) Hierarchical clustering of regBase Common integrated tools on three known functional/pathogenic regulatory variant datasets. HGMD, the Human Gene Mutation Database functional regulatory variants dataset; ClinVar, the ClinVar pathogenic and benign regulatory variants dataset; MPRA, the expression-modulating variants dataset identified by massively parallel reporter assay.

To visualize underlying relationships among these tools, we clustered the functional scores according to three above regulatory/pathogenic variant datasets. We found these tools could be generally partitioned into two major subsets, in which each member at the first subset barely associated with other tools within or outside this subset, while members at the second subset were usually correlated with each other (Figure 1B). This result indicates that some tools may capture the unique and important features that is able to distinguish regulatory variants from neutral ones. For example, deltaSVM and CATO learn classification models based on SNV disrupting DNase I hypersensitive site (DHS), and RSVP identifies many informative predictors from gene expression annotations. Interestingly, besides the tools that use exactly same training data or features, we found several tool pairs consistently clustered together in all three results, such as deltaSVM and CATO both utilize variants at DHS as training data. FATHMM-XF co-occured with CScape in the clustering, probably due to their use of similar negative samples and functional annotation features. (Figure 1B and Supplementary Figure S1). To summarize together, our results indicate that the existing noncoding variant functional scoring tools will produce inconsistent predictions across pathogenic/regulatory and neutral variants, and may capture various attributes of functional regulatory codes, suggesting the necessity and importance of systematic integration.

### Composite predictions of functional, pathogenic and cancer driver non-coding regulatory variant

Few ensemble prediction models for non-coding regulatory variants were proposed previously. These models only integrated limited number of tools and achieved mediocre performance on pathogenic regulatory variant prediction, especially for predicting somatic regulatory mutation associated with the development of cancer. Given the functional complexity and insufficient accumulation of causal regulatory variants, it is difficult to establish a well-rounded model that can predict all types of regulatory variants in the current stage. We hence partitioned the non-coding regulatory variant prediction task into three categories, including 1) predicting variant regulatory potential regardless of its functional direction and pathogenicity; 2) predicting disease-causal regulatory variant; 3) predicting cancer driver regulatory mutation. Correspondingly, we constructed three independent training datasets (Supplementary Table S6), including 1) functional regulatory variants dataset from our previous PRVCS (11) (regBase_REG); 2) pathogenic regulatory variants dataset from ClinVar and Genomiser (regBase_PAT); 3) highly recurrent regulatory somatic mutations dataset from COSMIC (regBase_CAN). For each positive set, we sampled constrained control set based on the best of our knowledge to alleviate biases (see Methods for details).

Owing to the potential complementarity and uniqueness of existing non-coding regulatory variant prediction algorithms, we hypothesized that combining functional scores from multiple tools would boost the prediction performance for each aforementioned regulatory variant category. Using the compiled golden standards and regBase scores, we trained three composite models by Gradient Tree Boosting (GTB). We adapted XGBoost classifier as the implementation of GTB algorithm (49), because sparsity-aware split finding of XGBoost make it suitable for the task as missing value are commonly appeared in our regBase features. As all training variants of regBase_REG came from 1000 Genomes Project, we were able to train additional model using regBase Common features (regBase_REG_Common). We tuned the model hyper-parameters by 10-fold cross-validation and evaluated the model performance by receiver operating characteristic (ROC) curve and area under the curve (AUC).

The new composite models significantly improved the prediction performance of the best single tool by 5% to 22% (Figure 2). Specifically, for functional non-coding regulatory variant prediction, regBase_REG_Common model received average AUC of 0.93 (Figure 2A) and regBase_REG model got 0.89 (Figure 2B). GenoCanyon is always the best single tool with AUC of 0.84 in these two models compared to an average score less than 0.75 achieved by the majority of tools, which implies that integrating more tools with weak but complementary ability could increase the performance of ensemble prediction model. For pathogenic non-coding regulatory variant prediction, regBase_PAT model reached an average AUC of 0.90 (Figure 2C) that exceeds the best tool ReMM by 6% (AUC of 0.84). Remarkably, Tools without training on any ClinVar data, like Eigen, LINSIGHT and CADD, can achieve a comparable performance (AUC > 0.8) with ReMM on predicting disease-causal regulatory variants. This may highlight that evolutionary information and unbiased leaning strategy frequently used in these tools, could be very useful to discriminate mutation pathogenicity or deleteriousness from neutral signals. For the prediction of cancer driver non-coding regulatory mutation, our regBase_CAN model got an unexpectedly high average AUC of 0.91 (Figure 2D) that outperformed the best tool FIRE by 22% (AUC of 0.69). We found most existing algorithms were not specially designed to prioritize somatic regulatory variants except for FunSeq2 and CScape in the regBase database. The preliminary understanding of regulatory codes in the cancer genome and the limited number of cancer driver non-coding variants could be keypoints that inhibited the development of effective prediction model. However, by compositing the effect of existing regulatory variant scoring scheme, we provided an alternative strategy to prioritize non-coding regulatory mutation with cancer driver potential.

**Figure 2.**
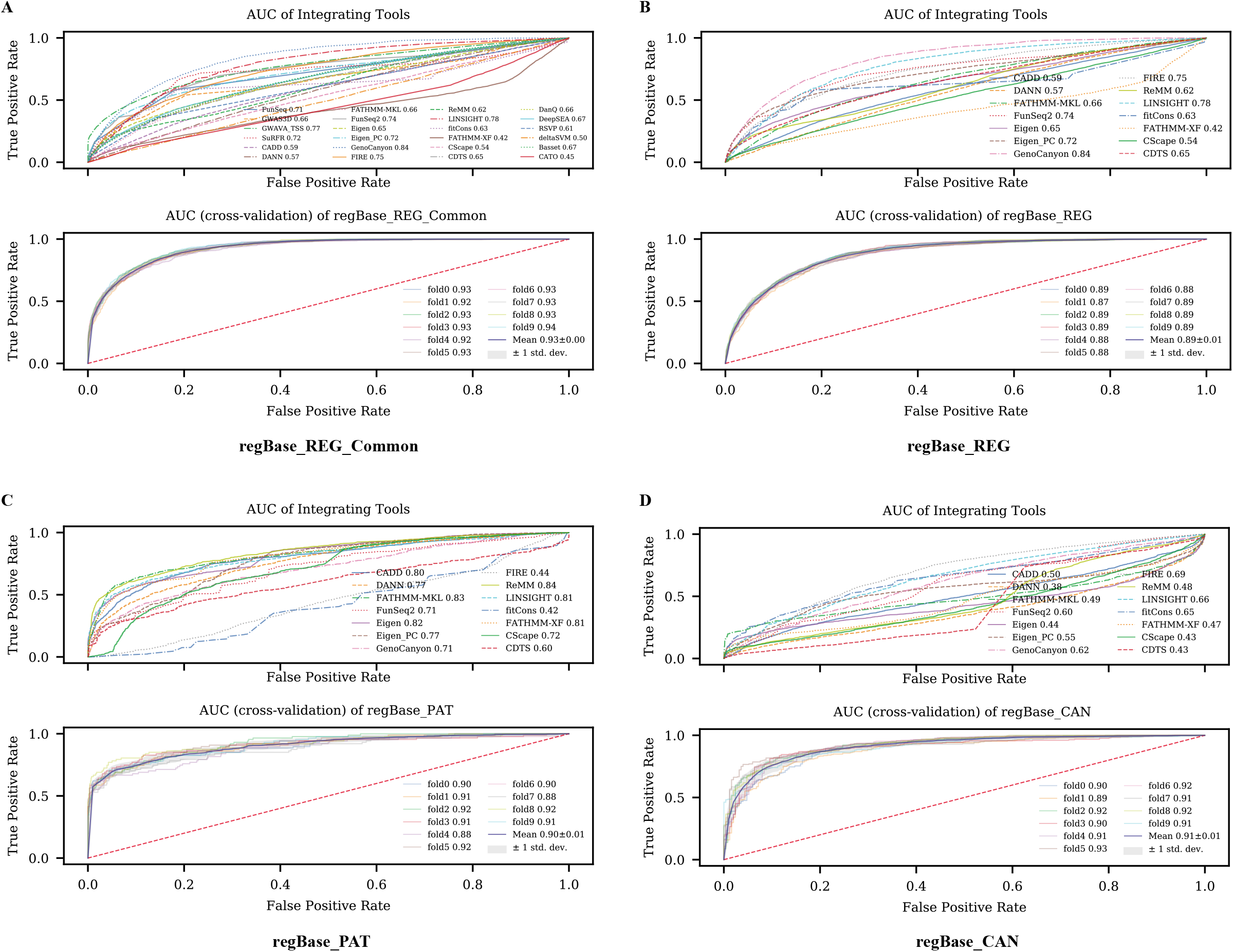
Receiver operating characteristic (ROC) curve and area under the receiver operating characteristics curve (AUC) for different prediction models using 10-fold cross validation. (A) ROC and AUC of 23 integrated tools and 10-fold cross validation result for regBase_REG_Common model. (B) ROC and AUC of 13 integrated tools and 10-fold cross validation result for regBase_REG model. (C) ROC and AUC of 13 integrated tools and 10-fold cross validation result for regBase_PAT model. (D) ROC and AUC of 13 integrated tools and 10-fold cross validation result for regBase_CAN model.

### Benchmarks on independent non-coding regulatory variant datasets

To systematically evaluate our four composite models, we constructed six independent benchmark datasets across different functional categories of non-coding regulatory variants (Supplementary Table S7), including two fine-mapped eQTL datasets (Brown_eQTL (50), GTEx_eQTL (52)), one experimental validated eQTL dataset (MPRA_eQTL (45)), two disease-associated variants datasets (GWAS_5E-8, GWAS_1E-5 (53)) and one somatic eQTL dataset (Somatic_eQTL (54)). We also sampled corresponding control testing dataset and removed variants that appeared in our training datasets. These independent datasets were not used to train almost all of integrated algorithms in the regBase database, which could provide an unbiased opportunity to comprehensively compare our models with existing tools.

In general, our composite models can achieve an AUC score around 0.8 for most of the above testing sets. Among them, regBase_REG_Common model was the best one to predict fine-mapped eQTLs (AUC of 0.88 for Brown_eQTL, AUC of 0.89 for GTEx_eQTL) and GWAS disease-associated SNVs (AUC of 0.88 for GWAS_5E-8, AUC of 0.83 for GWAS_1E-5) (Figure 3A), while the performance regBase_REG is similar but falls slightly behind (Figure 3B). This is consistent with the cross-validation results in model training step. For predicting expression-modulating variants identified by MPRA, the best composite model regBase_REG got relatively smaller AUC of 0.62, implying the integration of existing tools may have limited ability to distinguish sequence effect of transcriptional regulatory elements regardless of their chromatin context (Figure 3C). Regarding to the prediction of cancer relevant somatic eQTLs, regBase_CAN model received an AUC of 0.94 which largely outperformed other models, further indicating the combination of weak classifiers could generate stronger learner using Gradient Tree Boosting strategy (Figure 3D). Nevertheless, regBase_PAT model exhibited poor performance when predicting GWAS disease-associated variants. Compared with common germline variants that conferring hereditary disease predisposition, the pathogenic SNVs used to train regBase_PAT model are mostly rare variants to cause Mendelian disorders and obtain very distinct attributes. Therefore, other independent pathogenic dataset is needed to evaluate the actual performance of regBase_PAT model.

**Figure 3.**
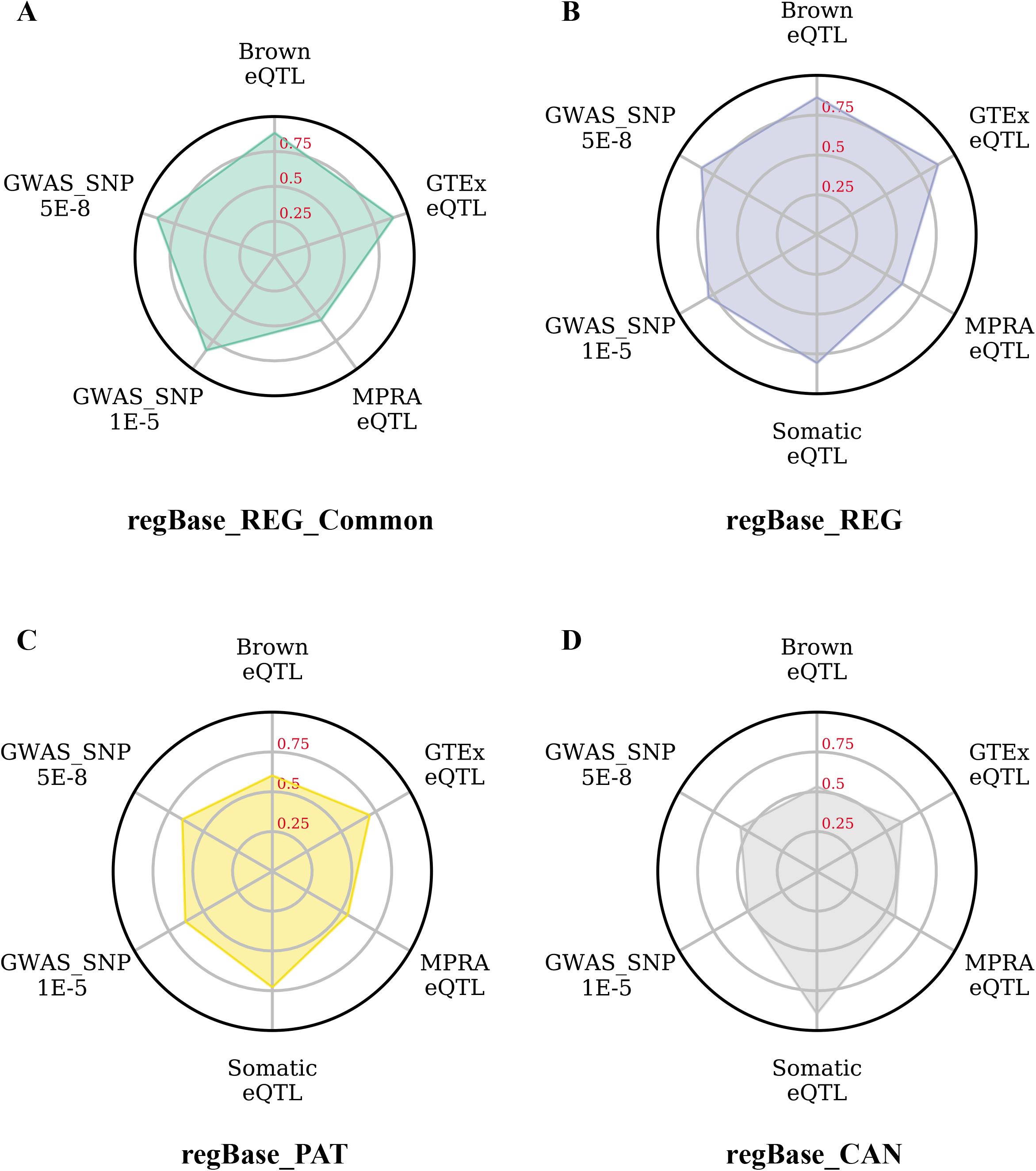
Area-under-curve scores distribution for six independent benchmarks. (A) regBase_REG_Common model. (B) regBase_REG model. (C) regBase_PAT model. (D) regBase_CAN model. Brown_eQTL, 11 tissue/cell type-specific eQTLs fine-mapping data that was profiled by Brown and colleagues; GTEx_eQTL, 44 tissues-specific eQTLs within fine-mapped credible set from GTEx V6; MPRA_eQTL, significant expression modulating variants by MPRA in lymphoblastoid cell lines; GWAS_5E-8, GWAS disease-associated variants with *P*-value < 5E-8 from GWAS Catalog; GWAS_1E-5, GWAS disease-associated variants with *P*-value < 1E-5 from GWAS Catalog; Somatic_eQTL, recurrent somatic mutations within significant flanking intervals per somatic eGene.

To figure out whether the combined models are better than individual tools or not, we evaluated the performance of 23 regBase Common integrated scores on five non-cancer testing sets, and 13 regBase integrated scores on somatic eQTL dataset. Results showed that our composite models outperformed individual tools on most of evaluations. First, regBase_REG_Common model was top ranked for Brown_eQTL (Figure 4A and Supplementary Table S8), GTEx_eQTL (Figure 4B and Supplementary Table S9), GWAS_5E-8 (Figure 4C and Supplementary Table S10) and GWAS_1E-5 (Figure 4D and Supplementary Table S11). It is worth noting that GenoCanyon, FIRE, LINSIGHT and Eigen_PC were well performed on predicting germline cis-eQTLs, while GenoCanyon, FunSeq2 and SuRFR were suitable to classify disease-associated regularity variants. In addition, regBase_CAN model was the best one for Somatic_eQTL dataset, with an AUC of 0.94 which greatly surpassed the second-best tool Eigen_PC (AUC of 0.86) (Figure 4E and Supplementary Table S12). Moreover, when predicting effective MPRA alleles, tools learned by deep learning or unsupervised model, such as DeepSEA, GenoCanyon, Eigen_PC and Basset, obtained a higher AUC than our regBase_REG model (Figure 4F and Supplementary Table S13), probably due to the fact that deep learning and unsupervised methods could capture unknown features that explain the *in-vitro* activity of regulatory allele. We also evaluated the performance of our newly trained models with existing ensemble methods including IW-Scoring (12) and our previous PRVCS (11). We found that regBase_REG_Common model obtained superior capability in five out of six benchmarks, except that PRVCS and IW-Scoring slightly outperformed regBase_REG model at MPRA_eQTL dataset (Supplementary Figure S2A-F and Supplementary Table S14). Taken together, these independent evaluations further demonstrated the effectiveness of our composite models and illuminated that non-coding regulatory variants prediction results could be increasingly applicable in the future genetic studies.

**Figure 4.**
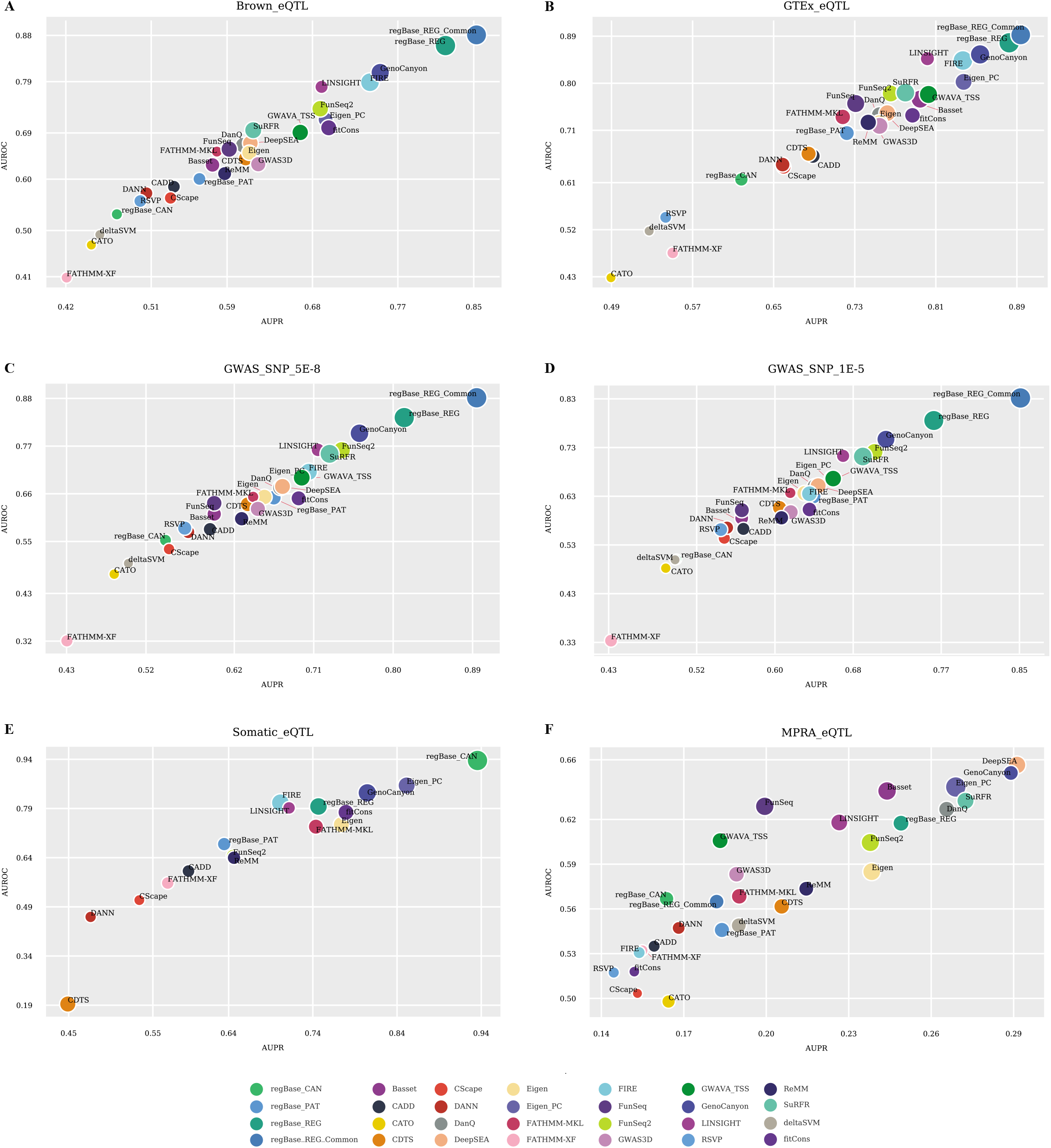
Evaluation result of individual prediction tools on six independent testing datasets. (A) Performance on Brown_eQTL dataset. (B) Performance on GTEx_eQTL dataset. (C) Performance on GWAS_5E-8 dataset. (D) Performance on GWAS_1E-5 dataset. (E) Performance on Somatic_eQTL dataset. (F) Performance on MPRA_eQTL dataset. AUPR, area under the precision recal curve; AUROC, area under the receiver operating characteristics curve; bubble size is proportional to Pearson correlation coefficients between predicted and true labels for each evaluation.

### regBase composite models facilitate the identification of causal non-coding regulatory variant from complex GWAS loci

Exploiting the true disease-causal variants is a challenging task in the GWAS study, especially for extremely high LD variants that locate in the non-coding genomic region. Statistical fine-mapping analysis usually ends with credible set of likely casual variants in which highly linked SNPs achieve similar posterior probabilities of causality, requiring further investigation of the true causal variants by other computational strategies, such as functional annotation (56). By visualizing regional PHRED-scaled score spectrum of composite models across 5p15.33 TERT region, we found several PHRED score peaks of regBase_REG, regBase_REG_Common and regBase_CAN generally colocalize with significant disease-associated variants identified by existing GWASs, especially in the TERT promoter region (Figure 5A and Supplementary Table S15). To evaluate the ability of our composite models for causal variant prioritization, we collected 22 unique SNPs in the 5p15.33 TERT region that confer risk of multiple cancers from ten GWAS fine-mapping results (Supplementary Table S16). Previous results showed there are many independent causal SNPs around the TERT genomic region, and many of them can alter promoter or enhancer activities (57). We revealed that our regBase_CAN and regBase_REG_Common models acquired relatively higher regional PHRED scores than other methods (tools with no more than 25% equal scores were selected) for collected fine-mapped SNPs (Figure 5B and Supplementary Table S17). Moreover, compared with relatively higher correlation among these 22 fine-mapped SNVs (Supplementary Figure S3), our top ranked variants (regional PHRED score > 10) of regBase_CAN or regBase_REG_Common showed very low LD with each other (Figure 5C), which indicates that our composite models could distinguish true signal from difficult credible set. For example, among all 22 prioritized fine-mapped SNPs by regBase_REG_Common model, rs2853669 obtained the largest PHRED score in the whole 5p15.33 TERT region (Figure 5C). This SNP was previously validated to disrupt TERT promoter and confer cancer risk by extensive functional experiments (58–60), further suggesting our composite model could efficiently narrow down the potentially causal variants for following functional validations.

**Figure 5.**
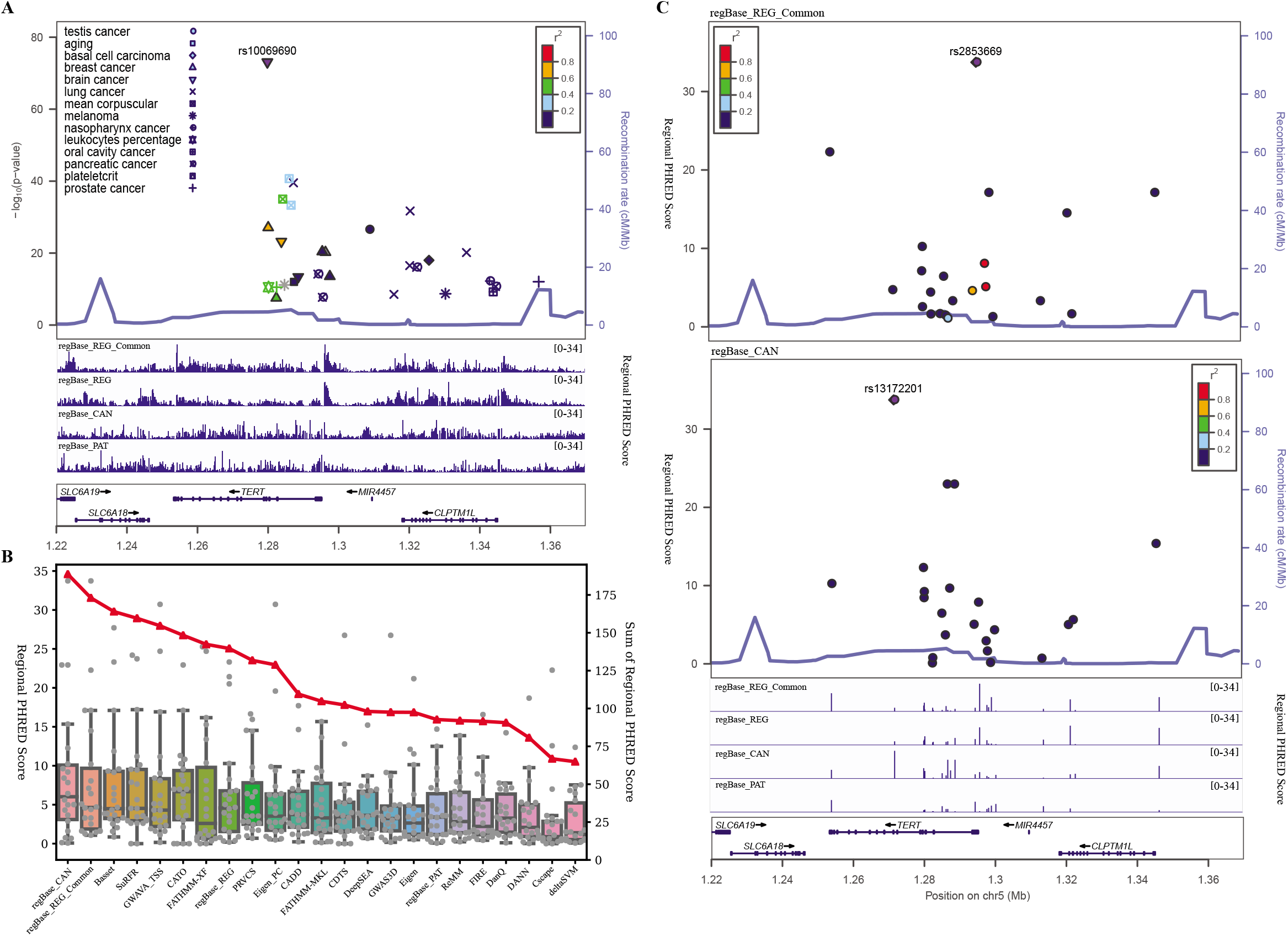
Non-coding regulatory variants prioritization at 5p15.33 TERT region. (A) GWAS significant SNPs and regional PHRED-scaled score distribution of our four composite models across 5p15.33 TERT region. LocusZoom plot is generated using the most significant SNP rs10069690 as lead and the EUR LD structure. (B) Comparison of regional PHRED scores among our composite models and all integrated methods for 22 fine-mapping SNPs at 5p15.33 TERT gene. Tools that obtain more than 25% equal scores in the evaluation are excluded. (C) LocusZoom plots for regional PHRED-scaled score of 22 fine-mapping SNPs. The top prioritized SNP rs2853669 in regBase_REG_Common model and the top prioritized SNP rs13172201 in regBase_CAN models are selected as leads.

## DISCUSSION

Evolved methods had been developed to predict and prioritize functional non-coding regulatory variants, yet systematical integration of existing predicted scores for all possible substitutions of human SNV was largely deficient. Comparing with a commonly used lightweight resource dbNSFP on functional prediction and annotation for human nonsynonymous and splice-site SNVs (61), we compile a comprehensive resource that includes 23 different tools to predict functional non-coding regulatory variants at the whole genome scale. To maximize the power and completeness for different types of non-coding regulatory variant prediction, we introduce three independent ensemble models to score functional, pathogenic or cancer driver regulatory variants respectively. We demonstrate that our composite strategies significantly increase the prediction accuracy and can greatly assist the casual noncoding regulatory variant discovery.

According to the benchmarks of several independent datasets, we found stable and reasonable performance of existing tools to predict variant regulatory potential regardless of its pathogenicity, such as predicting the probability of SNV to be a *cis*-eQTL. This merit could be attributed to the fact that current models are generally learned from annotation features that delineate regulatory signals around SNV locus, including chromatin accessibility, histone modifications and transcription factor binding. However, when evaluating the expression-modulating variants identified by *in vitro* reporter assay (62), no methods can achieve satisfactory performance. Since effective alleles in the MPRA are only weakly correlated with the associated eQTL effects (45,63), it may imply that surrounding sequence and local chromatin state could change the effect size of casual allele. In addition, recent CRISPR screening and GWAS fine mapping study have uncovered that some regulatory alleles locating in the unmarked regulatory elements are not associating with the conventional histone modifications or chromatin accessibility (64,65), which highlights the importance to exploit the missing but distinct prediction features. Besides, rational classification of pathogenic non-coding regulatory variant will extend the scopes of genetic diagnosis and precision medicine. Increasing studies have reported that pathogenic non-coding regulatory variant can influence the penetrance and causality of certain diseases (6), or alter the drug sensitivities (66,67). However, using ClinVar or COSMIC non-coding regulatory SNVs (not including splicing-altered SNVs) as golden standards (43,68), previous and our evaluations on pathogenic classification of regulatory variants showed limited performance (8,11). To this end, by leveraging the complementarity and uniqueness of existing methods, we trained regBase_PAT and regBase_CAN models to score the probability of variants being pathogenic or cancer driver in the gene regulation, and found significant improvements in both cross validation and independent benchmark. As the continual discoveries of non-coding disease-casual regulatory variants and more associated features, we believe that pathogenic prediction of non-coding regulatory variants will play a critical role in the clinical consensus interpretation of whole genome DNA sequence.

Highly context-dependent gene regulation can determine the cellular function of regulatory variants, and many recent methods are able to interpret regulatory variant in tissue/cell type-specific and disease-specific conditions (7,69). Since very few context-specific dataset could be used to benchmark the performance of tissue/cell type-specific predictions, researchers usually apply indirect solutions to evaluate the algorithms, such as the enrichment of tissue/cell type-specific epigenetic signals and *cis*-regulatory elements (70). Such imperfections and under calibrated performance could inhibit the broader applications of context-specific methods, especially for accurately predicting pathogenic regulatory variant on particular conditions. Despite the importance of systematic integration and evaluation of tissue/cell type-specific methods, regBase particularly aggregates and operates context-free prediction scores from existing tools. Our regBase aggregated scores together with three ensemble models provide a versatile tool that prioritizes organismal level non-coding regulatory variants in a context-free manner, greatly facilitating the interpretation of human non-coding genome in the era of precision medicine.

## DATA AVAILABILITY

The regBase models are implemented in Python. Integrated datasets, source codes, collected training/testing sets, analysis scripts for the results of this manuscript are available at https://github.com/mulinlab/regBase.

## FUNDING

This work was supported by grants from the National Natural Science Foundation of China 31701143, 31871327 (M.J.L.), Natural Science Foundation of Tianjin 18JCZDJC34700 (M.J.L.), and The Science & Technology Development Fund of Tianjin Education Commission for Higher Education 2018KJ082 (S.Z.).

## Conflict of interest statement

None declared.

